# Protease inhibitor ASP enhances freezing tolerance by inhibiting protein degradation in Kumquat

**DOI:** 10.1101/2022.10.17.512565

**Authors:** Hua Yang, Ke-wei Qiao, Jin-jing Teng, Jia-bei Chen, Ying-li Zhong, Li-qun Rao, Huang Li, Xing-yao Xiong

**Affiliations:** College of Bioscience and Biotechnology, Hunan Agricultural University, Changsha 410128, China; China Crop Germplasm Innovation and Resource Utilization, Hunan Agricultural University, Changsha 410128, China; Center for Plant Science Innovation, University of Nebraska-Lincoln, Lincoln, NE, 68588, USA

**Keywords:** Kumquat, cold acclimation, freezing tolerance, proteinase inhibitor, protein degradation

## Abstract

Cold acclimation is a complex biological process leading to the development of freezing tolerance in plants. In this study, we demonstrated that cold-induced expression of protease inhibitor FmASP in a citrus relative species kumquat (*Fortunella margarita* (Lour.) Swingle) contributes to its freezing tolerance by regulating protein degradation. First, we found that only cold-acclimated kumquat plants, although with extensive leaf cellular damage during freezing, are able to resume their normal growth upon stress relief. To dissect the impact of cold acclimation on this extraordinary freezing tolerance, we performed protein abundance assay and quantitative proteomics analysis of kumquat leaves subjected to cold acclimation (4 °C), freezing treatment (−10 °C) and post-stress recovery (25 °C). FmASP and a few non-specific proteases were identified as differentially expressed proteins induced by cold acclimation and associated with stable protein abundance throughout the course of freezing treatment. FmASP was further characterized as a robust inhibitor that inhibits the degradation capacity of multiple proteases. In addition, heterogeneous expression of *FmASP* in Arabidopsis confirmed its positive function in freezing tolerance. Finally, we proposed a working model of FmASP and illustrated how this extracellular-localized protease inhibitor protects proteins from degradation and consequently maintains essential cellular function for freezing stress recovery. These findings revealed the important role of protease inhibition on freezing response and provide insights on how this role may help develop new strategies to enhance plant freezing tolerance.

**One-sentence summary:** A protease inhibitor of *Fortunella margarita* enhances protein stability and freezing tolerance by regulating non-specific protease degradation

## Introduction

Freezing injury is a recurrent meteorological hazard that mainly affects overwintering crops, fruit trees, and economic forests (Rapacz et al., 2014; Ding et al., 2020). High-value horticultural crops are vulnerable to the threat of freezing temperature which has occurred more frequently in recent years due to climate change (Luedeling, 2012; Gupta and Deswal, 2014). To cope with and survive freezing temperatures, plants have evolved a series of cold responsive mechanisms which can be placed into two general categories: tolerance and avoidance (Gusta and Wisniewski, 2013). Freezing tolerance, defined as the ability of plants to survive extracellular freezing, is accomplished by the loss of cellular water to extracellular ice and the concomitant decease of freezing point in cytoplasm. It involves a cascade of transcriptomic and biochemical changes that frequently present in species found in locations where freezing events are severe and of long duration (Thomashow, 1999; Gusta and Wisniewski, 2013; Wisniewski et al., 2017). On the other hand, freezing avoidance mainly involves biophysical changes that regulate ice formation by allowing pockets of water to remain undercooled to very low temperatures, so that the supercooled cells are not exposed to the dehydrative effects and remain in a metastable condition. Deep supercooling feature of the avoidance mechanism is widely present in temperate tree species, such as in their xylem parenchyma cells and floral buds, from where the intensity of the freezing events is moderate and of short duration (George and Burke, 1984; Ashworth and Wisniewski, 1991; Wisniewski et al., 2014). Strictly speaking, freezing tolerance and avoidance are not mutually exclusive, as in both scenarios plants are trying to avoid or minimize cellular damage and hence improve their hardiness to freezing stress (Pearce, 2001; Gusta and Wisniewski, 2013). In the past two decades, with the advent of modern molecular biology and genetic resources, the cold signaling pathway and the underlying regulatory mechanisms of plant cold response has been extensively studied in model species Arabidopsis and staple crops (Chinnusamy et al., 2010; Kidokoro et al., 2017; Guo et al., 2018; Ding et al., 2019; Chang et al., 2021). Despite those advances, our understanding of cold acclimation in overwintering fruit crops, particularly its role in freezing stress recovery remains unclear.

Kumquat (*Fortunella margarita* (Lour.) Swingle) is a subtropical shrub widely cultivated for its rich nutrients and bioactive compounds. It is a close relative to citrus and commonly used in citrus fruit research and breeding (Khalaf et al., 2007). Compared to other deciduous citrus species, kumquat is evergreen and very tolerant to freezing temperatures (Smillie and Hetherington, 1983; Santini et al., 2013). In this study, we observed that cold-acclimated kumquat plants suffered from more than 50% damage rate of their leaves were still able to recover after being transferred to normal temperature condition. To investigate the underlying anti-freezing strategy, we carried out a global proteomic analysis of proteins that are responsive to cold acclimation and freezing treatment in kumquats. We identified several proteases and a protease inhibitor FmASP, specifically induced by cold acclimation, that contribute to kumquats’ freezing tolerance. Moreover, we experimentally confirmed that FmASP can effectively inhibit protein degradation and thus maintain protein stability during the course of freezing treatment, thereby retaining key cellular functions for freezing response and resistance.

## Results

### Cold acclimation enhances freezing tolerance of kumquat

We performed a freezing stress experiment on kumquat plants acclimated at 4 °C for two weeks (CA) with their non-acclimated control (NC) (stage 1). Without cold acclimation, kumquat leaves started to show frost damage around 1 h at -10 °C (stage 2) and suffered severe deformation and dehydration after 24 h recovery at 25 °C (stage 3). Permanent damage characterized by crisp texture and dark coloration was observed after recovery at 25 °C for one week (stage 4). In comparison, leaves from cold-acclimated kumquat plants exhibited minor changes in coloration during the treatment and can restore their color and shape in a week (Figure 1A). Similarly, the whole plants from the CA group resumed normal phenotype while the NC group suffered severe irreversible freezing damage and gradually wilted during the recovery period (Figure S1).

**Figure 1.**
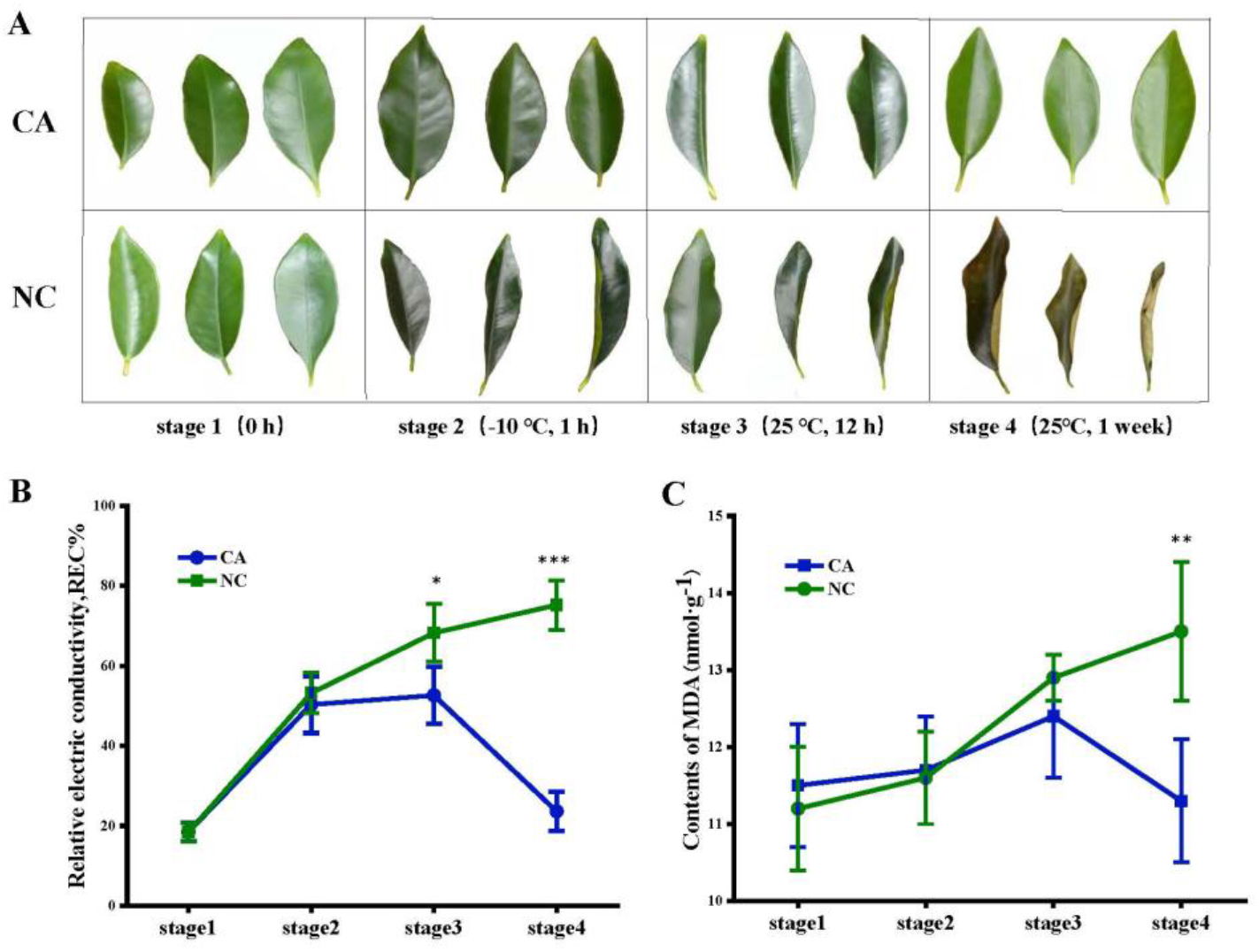
Cold acclimation confers freezing tolerance in kumquat plants. A, Phenotypic changes of cold-acclimated (CA) kumquat leaves subjected to freezing treatment with comparison to the non-acclimated control (NC). Stage 1, the end of cold acclimation; Stage 2, 1 h after -10 °C freezing treatment; Stage 3, 12 h recovery at 25 °C; Stage 4, one-week recovery at 25 °C. B, Measurement of the relative electrical conductivity (REC) of kumquat leaves from the CA group and the NC at four designated stages. C, Measurement of the MDA content of kumquat leaves from the CA group and the NC at four designated stages. Error bars indicate ± SE (n = 3). Asterisks indicate significant differences between the CA plants and the NC at the same stage (* P<0.05, ** P<0.01, *** P<0.001).

Membranes play a key role in enduring chilling injury in plant cell (Orvar et al., 2000; Moellering et al., 2010). Electrolyte leakage and MDA content are widely used as physiological indicators for membrane damage induced by various stresses (Tajvar et al., 2011; Jin et al., 2016). In this study, relative electric conductivity (REC) of kumquat leaves from the CA and NC group was quickly increased from 18.4% to 50.3% and 18.5% to 53.2% respectively at stage 2. REC of the CA and NC group continued to increase and reached 52.6% and 68.2% at stage 3. After recovery for one week (stage 4), REC of the control was 75.2% while REC of the CA group was decreased to 20%, close to the pre-treatment level (stage 1) (Figure 1B). Likewise, MDA content showed no significant difference between the CA group and the NC at stage 1 and 2. At the period of recovery, MDA content of the NC group exhibited a sharp increase at both stages (stage 3 and 4) while that of the CA group was increased first but then decreased to a level similar to stage 1 (Figure 1C). In consistent with the phenotypic observation, these results indicated that the extensive cellular damage caused by freezing can be restored in cold-acclimated kumquat plants.

### Cold acclimation maintains protein stability in kumquat leaves during freezing stress

Freezing stress is known to trigger a series of biochemical and physiological changes in plants at the protein level including the alteration of protein abundance and the production of cold-responsive proteins (Janmohammadi et al., 2015; Shi et al., 2018). In our protein content assay, we found that while the NC group exhibited significant losses of total protein on stage 3 and 4, the CA group maintained its protein content at a similar level throughout the treatment (Figure 2A). We observed a similar pattern of protein abundance change between the CA and NC group based on the gel band intensity in SDS-PAGE analysis (Figure 2B).

**Figure 2.**
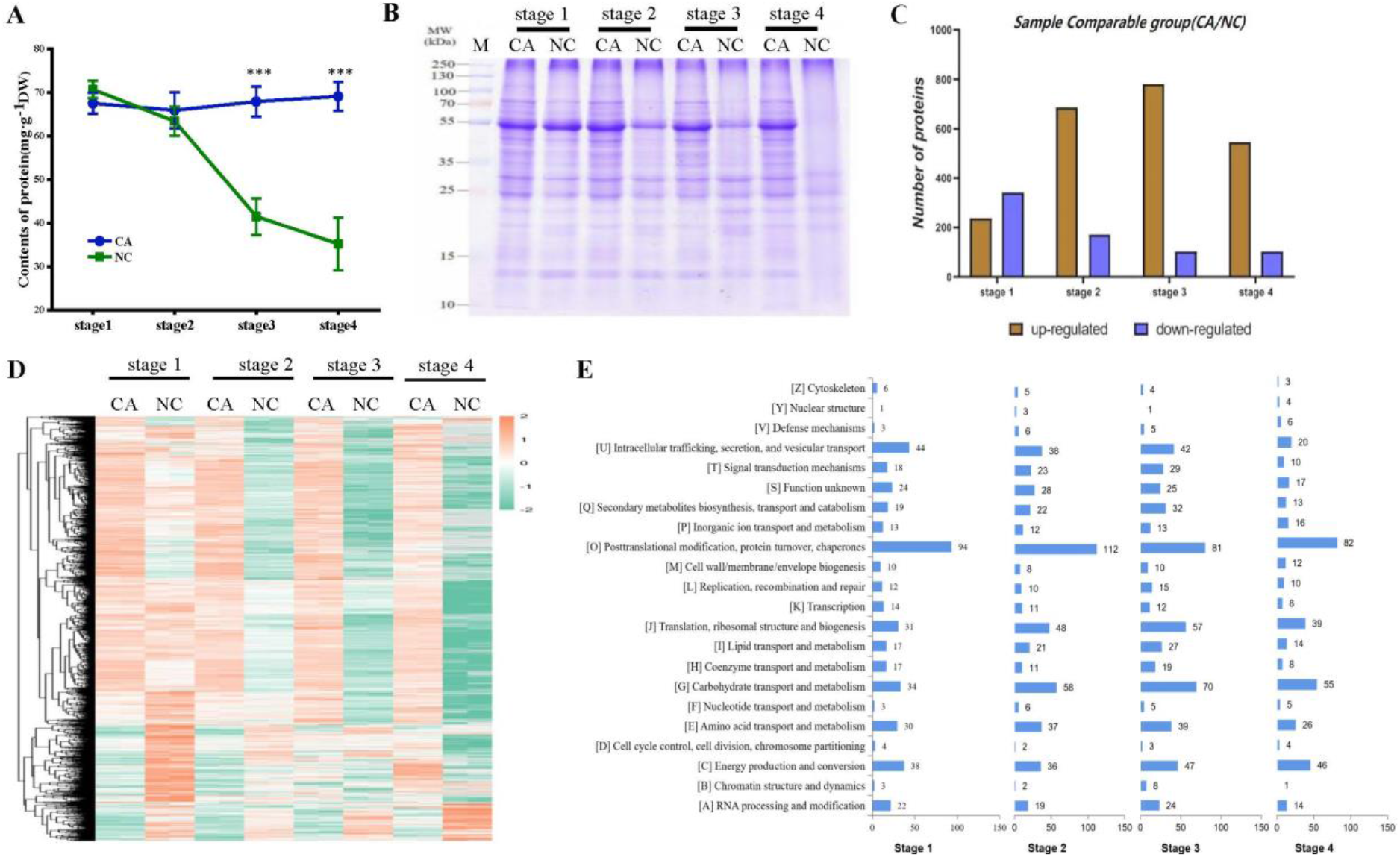
Cold acclimation contributes to the protein stability in kumquat leaves during freezing. A, Measurement of the protein content in kumquat leaves from the CA group and the NC at four designated stages. Error bars indicate ± SE (n = 3). Asterisks indicate significant differences between the CA plants and the NC at the same stage (*** P<0.001). B, SDS-PAGE analysis of total protein in kumquat leaves from the CA group and the NC at four designated stages. C, Number of up-regulated and down-regulated proteins in the CA group compared to the NC. D, Proteomic heat map of differentially expressed proteins between the CA group and the NC at four designated stages. E, Functional classification of differentially expressed proteins identified by proteomic analysis. Number of proteins in each category is shown on the right of each bar.

To identify differentially expressed proteins (DEPs) during the course of freezing treatment, we performed a global proteomics analysis on proteins extracted from kumquat leaf samples at all aforementioned stages. As a result, a total number of 3799 redundant proteins were confidently detected and quantified (Table S1). Using the same confidence level, we detected 580 DEPs that were induced by cold acclimation between the CA and NC group. We found surprisingly that the 1 h freezing resulted in a significant increase on the number of DEPs, especially the number of up-regulated proteins between the CA and NC at late stages (Figure 2C, Table S1). This distinct pattern indicated that 1) cold acclimation indeed impacted the freezing response of kumquat leaves at the proteomic level 2) the proportional change of upregulated protein abundance between CA and NC group could result from an increased de novo protein synthesis in CA or an increased protein degradation in NC or a combination of both. Thus, we next conducted clustering analysis to get an overview of individual protein changes based on normalized abundance at those four stages (Figure 2D). In general, the expression of a large number of clustered proteins from the NC group was reduced by freezing (stage 2) and continued the decreasing trend in the recovery phase (stage 3 and 4). By contrast, the expression of most proteins from the CA group remained constant in later stages (stage 3 and 4) after freezing treatment. More interestingly, some proteins downregulated by cold acclimation (stage 1) without any significant change when exposed to freezing stress can eventually restore their abundance during recovery (stage 4). Functional classification indicated that the DEPs were mostly involved in posttranslational modification, protein turnover, chaperons; intracellular trafficking, secretion and vesicular transport; energy production and conversion; and metabolic activities (Figure 2E). These results suggested that most of the acclimation-responsive proteins can either maintain or gradually retain their abundance while the non-acclimated group experiences significant fluctuation and loss of proteins during the course of freezing stress and recovery. Therefore, we concluded that cold acclimation is a critical process that confers kumquats’ freezing tolerance by maintaining protein abundance stability.

### Cold acclimation affects specific expression of protease and protease inhibitors

To further dissect the function of cold acclimation in freezing tolerance at proteomic level, we focused on the DEPs between the CA and NC group at stage 1. Differential protein abundance can be the result of several processes including changes in de novo protein biosynthesis, protein modification and protein degradation. Protease and protease inhibitors are common regulators for those changes during plant response to abiotic and biotic stresses (Thomas and van der Hoorn, 2018; Sharma and Gayen, 2021; Divekar et al., 2022). In this study, we identified a total of 31 cold-induced proteases mainly including the specific ATP-dependent Clp proteases, cysteine proteases and non-specific ones such as the serine proteases and aspartyl proteases (Table S2). Interestingly, the majority of non-specific proteases, located in chloroplast, were found to be down-regulated by cold acclimation. Moreover, we identified two differentially expressed protease inhibitors, ASP and KTI2 (Table S2). Notably, the upregulated ASP is homologous to the AtKTI5 (At1g17860.1), a member of the Kunitz trypsin inhibitor (KTI) family. KTI protease inhibitors have been found to be involved in plant immunity and stress resistance by regulating protease activity (Brzin and Kidric, 1996; Divekar et al., 2022). Taken together, we hypothesized that the cold-induced ASP could be one of a key players modulating freezing tolerance at the proteomic level in kumquat.

### FmASP acts as a major protease inhibitor in kumquat freezing response

To explore how FmASP functions as a protease inhibitor in the freezing response of kumquat, we studied its temporal expression, subcellular localization and inhibitory activity. In consistent with the proteomic results, FmASP is up-regulated by cold treatment (Figure 3A) and the FmASP protein is located in the extracellular space (Figure 3C), as revealed by qRT-PCR and the transient expression experiment, respectively. In addition, phylogenetic analysis revealed that FmASP is homologous to proteins characterized as the Kunitz trypsin inhibitors in multiple species including its closest citrus relative (Figure 3B). To test the inhibitory function of FmASP, we first expressed and purified the recombinant His-FmASP protein (Figure 3D), and then conducted an *in vitro* protein degradation assay. We found the addition of active FmASP protein can significantly reduce the degradation rate of soluble proteins extracted from kumquat leaves (Figure 3E). Specifically, its inhibitory effect can last up to 36 h to keep protein degradation at a much lower level compared to the control. Moreover, FmASP can inhibit the total proteolytic activity of common proteases and neutral protease as suggested in the inhibitory assay (Figure 3F). Among them, chymotrypsin and trypsin are the most effectively inhibited by FmASP, reflecting FmASP’s role as a Kunitz trypsin inhibitor.

**Figure 3.**
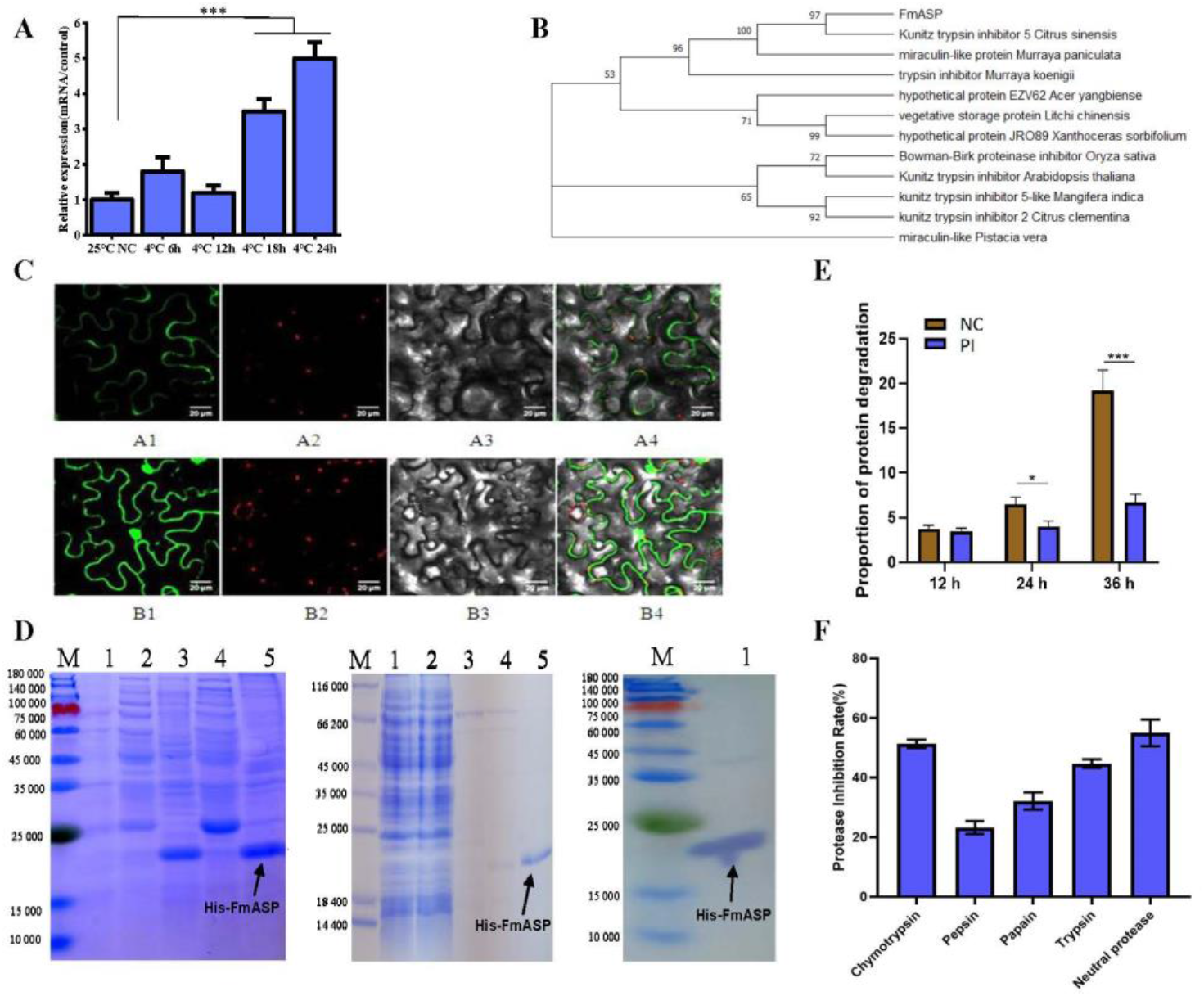
FmASP is an extracellular protease inhibitor in the KTI family. A, Relative gene expression of *FmASP* in the leaves of kumquat under cold stress as detected by qRT-PCR. Error bars indicate ± SE (n = 3). Asterisks indicate significant differences between the time points of treatment (*** P<0.001). B, Phylogenetic analysis of FmASP. Numbers above branches indicate bootstrap values. C, Subcellular localization of FmASP in tobacco cells revealed by the confocal microscopy. A1 represents the GFP fluorescence signal of the target gene; B1 indicates that the GFP fluorescence signal from an empty vector as the control; A2 and B2 indicate the chloroplast fluorescence signal; A3 and B3 represent the open bright field; A4 and B4 are the merged images of bright field, GFP and the chloroplast fluorescence. D, SDS–PAGE of the different steps in the purification of His-tagged FmASP protein and the Western blotting analysis. Left panel, Lanes: M, molecular weight marker; 1, total cell extract prior to IPTG induction; 2, soluble fraction of cells induced by IPTG at 20 °C; 3, pellet fraction of cells induced by IPTG at 20 °C; 4, soluble fraction of cells induced by IPTG at 37 °C; 5, pellet fraction of cells induced by IPTG at 37 °C. Middle panel, Lanes: M, molecular weight marker; 1, cleared elute of His-tagged protein extract; 2, flow through from the HisTrap column; 3. elute of the washing buffer with 20mM Imidazole; 4. elute of the washing buffer with 50mM Imidazole; 5. elute of the final wash with 500mM Imidazole. Right panel, Lanes: M, molecular weight marker; 1. the target protein was detected on Western blotting with an anti-His-tag antibody. Arrows indicate the positions of His-FmASP protein (21 KDa). E, *In vitro* protein degradation assay using FmASP as the protease inhibitor (PI). F, *In vitro* inhibitory assay on serve proteases using FmASP.

### Heterogeneous expression of *FmASP* enhances freezing tolerance in Arabidopsis

To further confirm the protease inhibitory function of FmASP in freezing stress response, we generated stable transgenic Arabidopsis plants overexpressing the *FmASP* gene and subjected them to freezing tolerance assay. We also generated independent *FmASP*-silencing Arabidopsis plants by RNAi in the background of the overexpressed line for comparison. Compared to the wild type, overexpression of *FmASP* significantly enhanced freezing tolerance as suggested by higher survival rate at the recovery stage (Figure 4A, 4B, 4C). Measurements of their electrical conductivity and MDA content further indicated that the membrane damage incurred by freezing can be restored in *FmAS*P-overexpressing plants (Figure 4D, 4E). In addition, an assay of total protein content revealed that *FmASP*-overexpression lines can maintain their protein content at a more constant level after exposing to freezing stress, as compared to that of the wild type (Figure 4F). The *FmASP*-silencing plants exhibited an intermediate survival rate and substantial cellular damage similar to that of the wild type (Figure 4A-F). These results implied that *FmASP* can enhance freezing tolerance in Arabidopsis by maintaining the overall protein stability and largely restoring the cellular injury to a pre-freezing level, whereby plants are able to resume growth.

**Figure 4.**
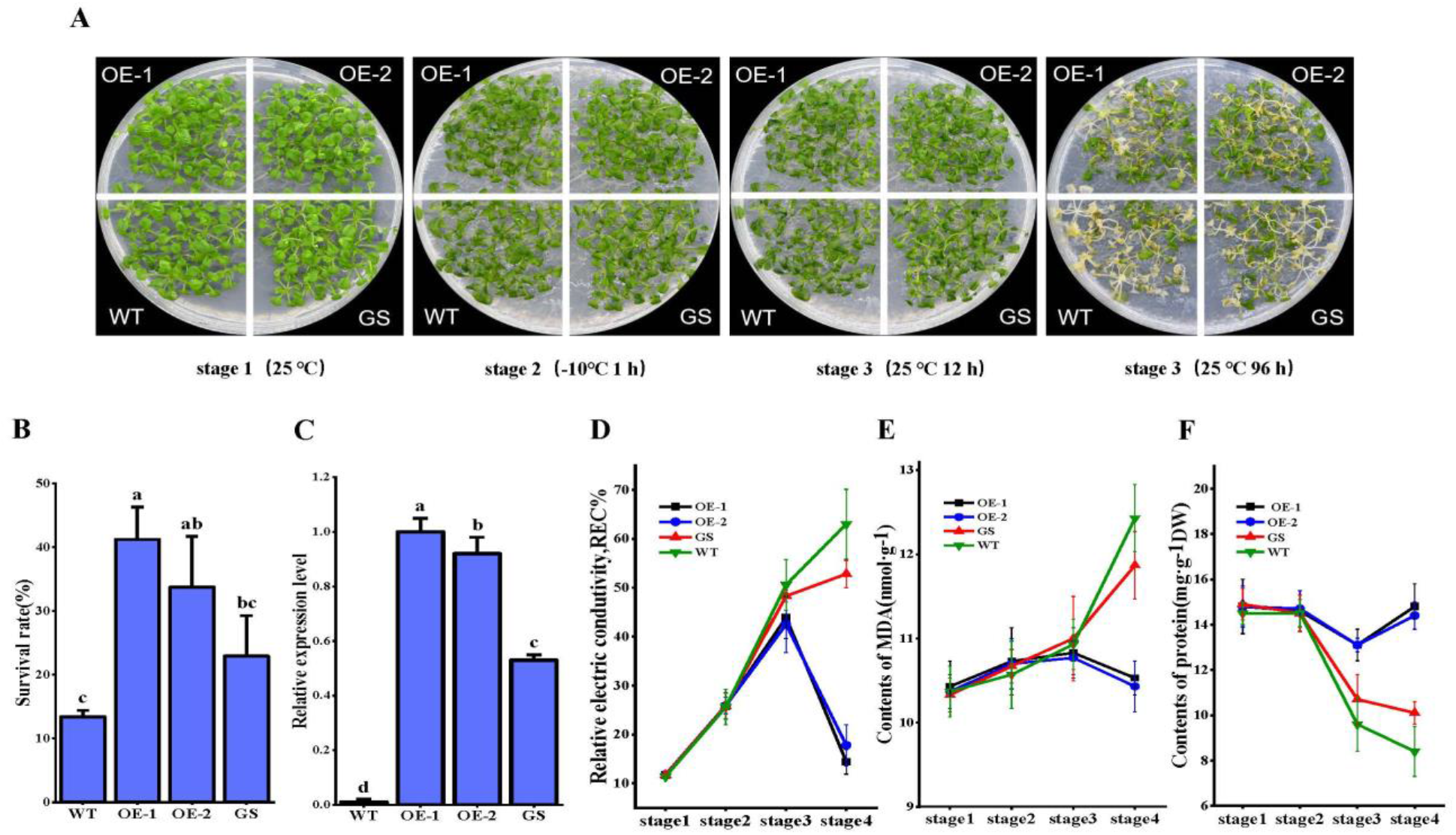
Overexpression of *FmASP* confers freezing tolerance in transgenic Arabidopsis plants. A, Plant phenotypes of the transgenic Arabidopsis lines and WT before and after the cold treatment. OE-1, overexpression line 1; OE-2, overexpression line 2; GS, gene-silencing line; WT, wild type. Stage 1: three-week-old seedlings grown at 25 °C; Stage 2: 1 h after -10 °C freezing treatment; Stage 3: 12 h recovery at 25 °C; Stage 4: 96 h recovery at 25 °C. B, Survival rates were counted at stage 4. Data are shown as means of three replicates ± SD. Different letters on the top of each bar denote statistically significant differences among the lines at P < 0.05 level. C, Relative gene expression of *FmASP* in the transgenic plants and WT as detected by qRT-PCR. D, Measurement of the relative electrical conductivity (REC) in the transgenic plants and WT at all four stages. E, Measurement of the content of MDA in the transgenic plants and WT at all four stages. F, Measurement of the protein content in the transgenic plants and WT at all designated stages.

## Discussion

Plant cold acclimation is very complex, involving a series of inducible mechanisms to protect cells from freezing injury (Thomashow, 1999; Xin and Browse, 2000). Among those cryoprotective mechanisms, protein stabilization plays a crucial role in maintaining the structure and function of delicate cell membranes during freezing stress (Crosatti et al., 2013; Chang et al., 2021). Plant proteases are ubiquitous enzymes that play an essential role in regulating protein quality and homeostasis to acclimatize environmental stresses (van der Hoorn, 2008; Sharma and Gayen, 2021). Several types of proteases have been characterized to be associated with plant responses to drought (Yao et al., 2012; le Roux et al., 2019), salinity(Jones and Mullet, 1995; Chen et al., 2010; Mishra et al., 2016) and biotic stresses(Hatsugai et al., 2004; Thomas and van der Hoorn, 2018). In this study, we investigated the temporal changes of protein abundance in kumquat plants under a regime of cold acclimation, freezing exposure and stress recovery. We found cold-acclimated plants had a better control on protein content stability during the whole treatment regime. Using label-free quantitative proteomics, we identified a total of 31 proteases significantly induced by cold acclimation, the key step determining whether a kumquat plant can recover from the severe freezing injury indicated by membrane ion leakage and oxidative damage. The differentially expressed proteases are distributed across different cellular compartments, especially on the plastid membranes (Table S2). They mainly include ATP-dependent Clp (serine-type) proteases and cysteine proteases, two protease families with potential implications in plant stress responses (Chen et al., 2010; Mishra et al., 2016). Specifically, Clp proteases are a prominent protease family located in the chloroplast that degrades numerous stromal proteins (Adam et al., 2006). Cysteine proteases are the best characterized protein proteases involved in many processes, particularly those associated with storage protein degradation and programmed cell death (Grudkowska and Zagdanska, 2004; Muntz, 2007). Taken together, we concluded that the induction of the Clp, cysteine proteases and other non-specific proteases during cold acclimation could be one of the main factors contributing to protein stability and thus the freezing tolerance in kumquat.

Moreover, we identified a cold-induced protease inhibitor FmASP in proteomics study and characterized its function. Protease inhibitor are multifunctional proteins implicated in the control of endogenous proteolysis under biotic and abiotic stress conditions (Brzin and Kidric, 1996; Divekar et al., 2022). Constitutive expression of protease inhibitors has been shown to enhance the stress response and tolerance in plants (Huang et al., 2007; Srinivasan et al., 2009; Tiwari et al., 2015). However, the involvement of protease inhibitors in plant cold response and the underlying mechanism are largely unknown. In agreement with the positive role of protease inhibitors in plant immune and stress responses, we demonstrate that FmASP, as a member of the Kunitz trypsin inhibitor (KTI) family, confers the freezing tolerance in kumquat and Arabidopsis plants. In vitro protein degradation and enzyme activity assays on purified FmASP protein confirmed its role as a protease inhibitor. We also observed a gradually increased expression pattern of FmASP at 4 °C and that the inhibitory activity of FmASP can last for at least 36 h, reflecting its role as a plant defensive chemical in natural conditions. Overexpression of *FmASP* enhanced the freezing tolerance in Arabidopsis, whereas *FmASP*-silenced transgenic lines were more susceptible to freezing damage. Notably, the degree of cellular damage indicated by ion leakage, MDA quantification and the soluble protein content were positively associated with the FmASP expression level during freezing stress. Our results demonstrated that cold-induced FmASP could inhibit proteases and thus the overall stability of vital cellular proteins increases leading to enhanced freezing tolerance.

Subcellular localization showed FmASP is an extracellular protease inhibitor. An intriguing question is how FmASP enters the cells from the apoplast during cold acclimation, as this seems to be an induced but highly regulated process. Several plant proteases and protease inhibitors have been found to enter the cells predominately through endocytosis, a sophisticated mechanism which can avoid the unwanted presence of these proteolytic enzymes in cytosol to degrade essential constituents (Cui et al., 2013; Trusova et al., 2019; Trusova et al., 2019). In our proteomic analysis, we did find multiple proteins involved in the endocytic pathway are upregulated by cold stress (Table S3). Further investigation of the activation mechanism of endocytosis and the trafficking of FmASP will help us to understand more about this understudied area of plant cold acclimation.

Based on all current results, we propose a new strategy of plant resilience with the involvement of proteases and protease inhibitors that confers the freezing tolerance in kumquat (Figure 5). During cold acclimation, non-specific proteases from different cellular compartments are downregulated, whereas the extracellular-localized FmASP is upregulated. In addition, the rearrangement of plasma membrane components is initiated and probably accelerated by endocytosis-related proteins which are induced under cold conditions (Table S3). FmASP could enter the cells in the form of endocytic vesicles during the process of endocytosis. Upon freezing exposure, the gradual formation of cellular ice causes damage in membranes including that of endocytic vesicles. Then, FmASP is released to the cytoplasm and in close contact with the proteases. And their enzymatic activities are mostly preserved at freezing temperatures. Upon rewarming, FmASP acts its inhibitory ability on proteases and prevents nonspecific protein degradation. As a result, most of the functional proteins including cold-responsive proteins in the cells avoid degradation and resume their function to cope with freezing stress, thereby enhancing the freezing tolerance in cold-acclimated kumquat. This working model of protease inhibitor FmASP will be of great importance for improving our understanding of freezing tolerance of plants and provide a new perspective to manipulate the responsive mechanisms of environmental stresses in temperate fruit tree species.

**Figure 5.**
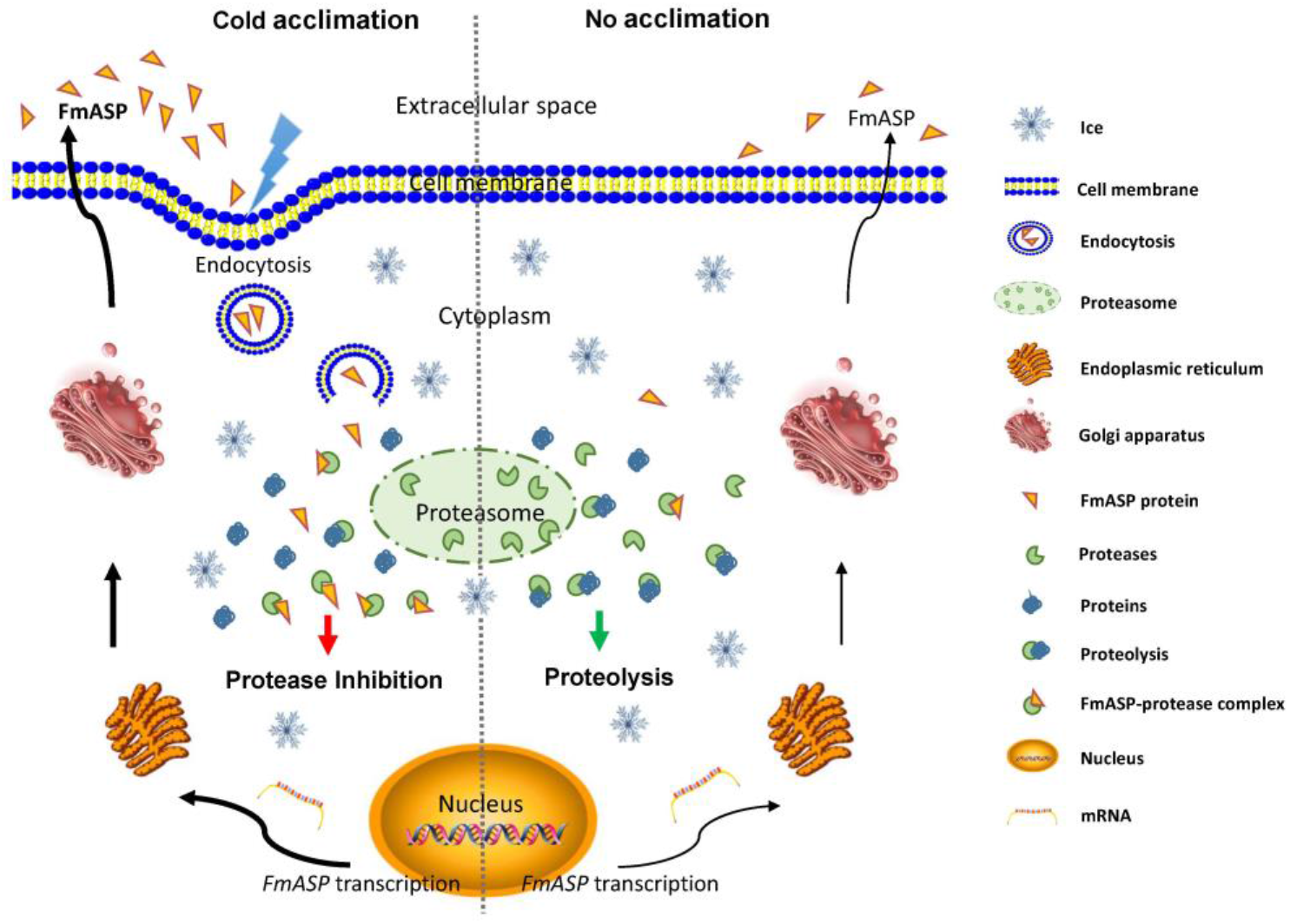
A working model of FmASP in freezing tolerance.

Compared to the non-acclimated condition, cold acclimation induces the expression of the nuclear-encoded gene *FmASP* and the concomitant protein production and extracellular transport. With the proceeding of cold stress, the rearrangement of cell membrane components is activated. FmASP protein accumulated in the extracellular space could enter the cell in the form of endocytic vesicles during the process of endocytosis. Upon freezing exposure, ice formation causes damage in membranes including that of endocytic vesicles. Then, FmASP is released to the cytoplasm and in close contact with proteases. And their enzymatic activities are mostly preserved at freezing temperatures. Upon rewarming, FmASP acts its inhibitory ability on proteases and thus prevents protein degradation. As a result, most of the functional proteins including the cold-responsive proteins avoid degradation and resume their function to cope with freezing stress, thereby enhancing freezing tolerance.

## Materials and methods

### Phenotypic observation of kumquat plants under freezing stress

Six-month-old kumquat (*Fortunella margarita* Swingle) seedlings from the National Center for Citrus Improvement (Changsha, China) were transplanted to pots with the compost soil (potting soil, turfy soil, and vermiculite 1:1:1) and grown for about 1.5 years in a greenhouse, with ambient temperature between 25 and 30 °C. We selected twenty well-established plants of similar size for the following treatment regime. Ten plants were transferred to a climate-controlled growth chamber for cold acclimation (CA) group at 4 °C while the other ten plants were grown in parallel at 25 °C as the non-acclimated control (NC) group (stage 1). After one month, all plants were placed in a freezing chamber (-10 °C) for 1 h (stage 2) and then transferred to the growth chamber for recovery. The plants recovered in the chamber at 25 °C for 24 h and for one week were recorded as stage 3 and 4, respectively. Three independent freezing tolerance assays were performed as replicates. Representative leaves and whole plants from each stage were selected for photographing. The sample leaves were collected at those four designated stages, flash frozen in liquid nitrogen for subsequent analysis.

### Measurement of physiological indicators in freezing-stressed leaf tissue

Relative electric conductivity (REC) was measured using a Mettler Toledo FE30 conductivity meter. Kumquat leaf samples were placed into 20 mL tubes containing 10 mL deionized water. The solution was vacuumed for 30 min and the conductance of water was measured as S0. After shaken at room temperature for 1 h, the conductance of water was measured as S1. Then samples were boiled in boiling water for 30 min and shaken at room temperature for 1 h with the conductance measured as S2. The value of (S1-S0)/(S2-S0) was calculated as the REC.

Malondialdehyde (MDA) content was determined by spectrophotometry following a protocol (Yang et al., 2011) with slight modification. Briefly, fresh kumquat leaves (0.5 g) were ground with a mortar and pestle and homogenized in 1 mL of 10% trichloroacetic acid (W/V). The homogenate was transferred to a centrifuge tube, washed with 1 mL 10% trichloroacetic acid, and centrifuged for 10 min at 10000 g. Next, 1 mL of supernatant was mixed with 1 mL of 0.6% thiobarbituric acid and incubated in boiling water for 30 min. After brief cooling on ice, the solution was centrifuged at 3000 g for 10 min, and the absorbance of the supernatant was measured at 450, 532 and 600 nm respectively. The content was calculated as MDA (nmol.g^-1^) = 6.45 × (A532 − A600) − 0.56 × A450.

The soluble protein content of kumquat leaves from each designated stage was measured by the coomassie brilliant blue method.

### Protein preparation, digestion and LC-MS/MS analysis

Approximately 500 mg leaves sampled from the CA or NC group at each stage was ground thoroughly in liquid nitrogen and extracted with 5 mL 1% SDS lysis buffer containing 10 mM dithiothreitol and 1% protease inhibitor. An equal volume of Tris-saturated phenol (pH 8.0) was added before the mixture was vortexed for 5 min. After centrifugation (4 °C, 10 min, 5500 g), the upper phenol phase was transferred to a new centrifuge tube. Proteins were precipitated by adding five volumes of ammonium sulfate-saturated methanol and incubated at −20 °C overnight. The produced precipitate after centrifuge was washed with cold methanol, followed by three washes with acetone. Protein concentration was determined using a bicinchoninic acid (BCA) protein assay kit (Beyotime, China). For digestion, equal amount of protein from each sample was adjusted to the same volume using lysis buffer and then re-precipitated with 20% TCA at 4 °C for 2 h. After washing twice with ice-cold acetone, the precipitate was diluted in 200 mM tetraethylammonium bromide (TEAB, pH 8.0). Finally, trypsin was added for digestion (w/w for enzyme: sample = 1: 50) at 37 °C for 16 h. The resulting peptide mixture was reduced with 5 mM dithiothreitol (DTT) for 30 min at 56 °C and alkylated with 11 mM iodoacetamide for 15 min at room temperature in darkness.

Proteomic data acquisition was performed on an EASY-nLC 1200 ultra-performance liquid chromatography system connected to an Orbitrap Exploris 480 mass spectrometer (Thermo Scientific, San Jose, CA). For LC-MS/MS proteomic data analysis, we used Maxquant (v1.6.15.0) with the search database (Citrus_japonica_76966_TX_20210429.fasta) to process raw MS files. Search results were filtered at 1% false discovery rate (FDR) and peptide confidence level was set as at least one unique peptide per protein for protein identification. The up- or down-regulated proteins with relative quantification *p*-values < 0.05 and 1.5 fold-change ratios were selected as differentially expressed proteins. Euclidean distance and hierarchical cluster were used to cluster differentially expressed proteins. Functional annotation and the analysis of annotation data were performed using Blast2GO (https://www.blast2go.com/). KEGG database (http://www.genome.jp/kegg/) and Clusters of Orthologous Groups of proteins (COGs) (http://www.ncbi.nlm.nih.gov/COG/) were used for protein identification, classification, and clustering.

### Subcellular localization, heterologous expression and purification of FmASP

Full-length cDNA of *FmASP* fused with a C-terminal green fluorescent protein (GFP) was cloned into vector pCAMBIA2300. The constructed plasmids were transferred into Agrobacterium strain *EHA105* and then transformed into tobacco (*Nicotiana benthamiana*) leaves for transient expression. Subcellular localization of GFP fusion proteins was detected and captured by a laser confocal microscope (FV1000-IX81, Olympus, Japan).

Recombinant plasmid pBRT7-7806 and the control plasmid pBRT7 were synthesized and transformed into E. coli BL21 (DE3) to express FmASP protein. Target protein was recovered and purified with a His-Trap affinity column (QIAGEN) according to manufacturers’ protocols. His-FmASP protein was detected by immunoblotting with using a mouse anti-His antibody (GenScript, A00186). The immunoblotting band signals were visualized by enhanced enhanced chemiluminescence (ECL) detection system (Tanon, Shanghai, China) and images were obtained using a cooled charge-coupled device (CCD) camera (Tanon-4100).

### Proteinase inhibition assays

Kumquat leaves were pulverized in liquid nitrogen using a pestle and mortar, weighed and incubated on ice with 10 mL PBS buffer (pH 6.5) for 30 min with occasional mixing. The mixture was centrifuged at 20000 g at 4 °C for 30 min and the resulting supernatant was centrifuged two more times. 0.9 mL of the supernatant was placed in a test tube with the addition of 1 μl purified FmASP protein. Total protein content in the solution after incubating at 25 °C for 0, 12, 24 and 36 h was measured using the BCA protein assay kit (Beyotime, China). Protease inhibition was assessed as the percentage of reduction in protein concentration relative to that of the control. Control for each set was made by adding equal volume of heat-denatured FmASP. All assays were made with three different pools and in triplicate. The inhibitory activity of FmASP was measured in PBS buffer containing 20μg/ml protease cocktail (chymotrypain, pepsin, papain, trypsin and neutralprotease) with 200μg/ml S-7388 substrate (Sigma-Aldrich); final volume was 100 μL. Prior to substrate addition, proteases were incubated for 30 min at 37 °C with either 10μg/ml FmASP or equal volume of PBS buffer. The reaction velocity was measured spectrophotometrically at 450 nm for 5 min using a SpectraMax M5e plate reader (Molecular Devices, USA). Experiments were performed in triplicates and velocities were reported as means ± SD.

### Plasmid construction and generation of transgenic plants

Full-length ASP cDNA obtained from *F. margarita* was amplified by RT-PCR using primers with compatible enzyme digestion sites. The *FmASP* amplicon was confirmed by sequencing and then cloned into a p1300M vector under control of the constitutive cauliflower mosaic virus 35S promoter. The 35S::p1300M-ASP construct was introduced into the *Agrobacterium* strain GV3101 and then transferred into wild-type Arabidopsis (Col-0) plants by floral-dip transformation. Transgenic lines obtained were first screened by hygromycin and then verified by PCR. T4 homozygous transgenic lines were selected for freezing tolerance assay and downstream analysis.

Gene silence vector pBWA(V)HS-ASP was constructed by assembling inverted repeats of 200-300 bp fragments of *FmASP* CDS linked by a 200 bp loop sequence and driven by the CaMV 35S promoter. After verified by DNA sequencing, the vector was transformed into the homozygous FmASP overexpression line of Arabidopsis to obtain gene silencing (GS) lines. The information of all primers used in this study was presented in Table S4.

### qRT-PCR assay

Total RNA was extracted from 4-week-old Arabidopsis plants with TRIzol reagent (Invitrogen). Reverse transcription and quantitative real-time PCR assay were performed in a Bio-Rad iQ5 real-time system using a Quant One Step qRT-PCR (SYBR Green I) Kit (Tiangen Biotech). Transcript expression levels of *FmASP* were normalized to that of the reference gene *ACTIN* and calculated using the 2^-ΔΔCt^ method (Livak and Schmittgen, 2001).

### Phenotypic observation, physiological assays of transgenic Arabidopsis under freezing stress

Three-week-old seedlings grown on ½ MS medium plates from a climate-controlled growth chamber (25 °C; stage 1) were placed into a freezing chamber at -10 °C for 1 h (stage 2) and then recovered at 25 °C for 12 h (stage 2) and 96 h (stage 3) before counting the survival rate. Seedlings with a non-dehydrated stem and at least three green leaves were recorded as the survivors. The MDA, relative electric conductivity and total protein content assays of transgenic Arabidopsis were performed using the same methods as described in kumquat experiments.

## Supplemental Data

**Supplemental Figure S1**. Phenotypic evaluation of non-acclimated and cold-acclimated kumquat plants under freezing stress.

**Supplemental Table S1**. Summary and list of differentially expressed proteins and peptides in proteomic analysis.

**Supplemental Table S2**. List of proteases and protease inhibitors identified as differentially expressed proteins induced by cold acclimation.

**Supplemental Table S3**. List of differentially expressed proteins associated with endocytosis in cold acclimation.

**Supplemental Table S4**. List of primers used in this study.

## Acknowledgments

The authors would like to thank Drs. Xiangyang Lu, Zhanguo Xin and Huazhong Shi for giving technical guidance and Drs. Zhanguo Xin and Xinbo Chen for the revision of this paper.

## Author contributions

H.Y., X.X. and H.L. conceived and designed the experiments. H.Y., K.Q., J. T., J.C., and Y.Z. set up and carried out the experiments. H.Y., K.Q., J.T., Y.Z., L.R. and H.L. analyzed the data. H.Y., Y.Z. and H.L. wrote the paper. All authors reviewed the manuscript.

